# High Dimensional Mediation Analysis with Applications to Causal Gene Identification

**DOI:** 10.1101/497826

**Authors:** Qi Zhang

## Abstract

Mediation analysis has been a popular framework for elucidating the mediating mechanism of the exposure effect on the outcome. Previous literature in causal mediation primarily focused on the classical settings with univariate exposure and univariate mediator, with recent growing interests in high dimensional mediator. In this paper, we study the mediation model with high dimensional exposure and high dimensional mediator, and introduce two procedures for mediator selection, MedFix and MedMix. MedFix is our new application of adaptive lasso with one additional tuning parameter. MedMix is a novel mediation model based on high dimensional linear mixed model, for which we also develop a new variable selection algorithm. Our study is motivated by the causal gene identification problem, where causal genes are defined as the genes that mediate the genetic effect. For this problem, the genetic variants are the high dimensional exposure, the gene expressions the high dimensional mediator, and the phenotype of interest the outcome. We evaluate the proposed methods using a mouse f2 dataset for diabetes study, and extensive real data driven simulations. We show that the mixed model based approach leads to higher accuracy in mediator selection and mediation effect size estimation, and is more reproducible across independent measurements of the response and more robust against model misspecification. The source R code will be made available on Github https://github.com/QiZhangStat/highMed upon the publication of this paper.

## 1 Introduction

Mediation analysis is a type of causal inference investigating the effect of certain exposures (*Z*) on an outcome (*Y*, Figure 1a). It assumes a hypothesized causal chain in which all or a part of the exposure effect on the outcome may be attributed to its effect on some mediators (M) which in turns influence the outcome (indirect effect). These variables could be also influenced by some observed confounders (*X*). Mediation analysis was originated in social science (Baron and Kenny, 1986; MacKinnon, 2012), and has been widely used in many applications.

**Figure 1:**
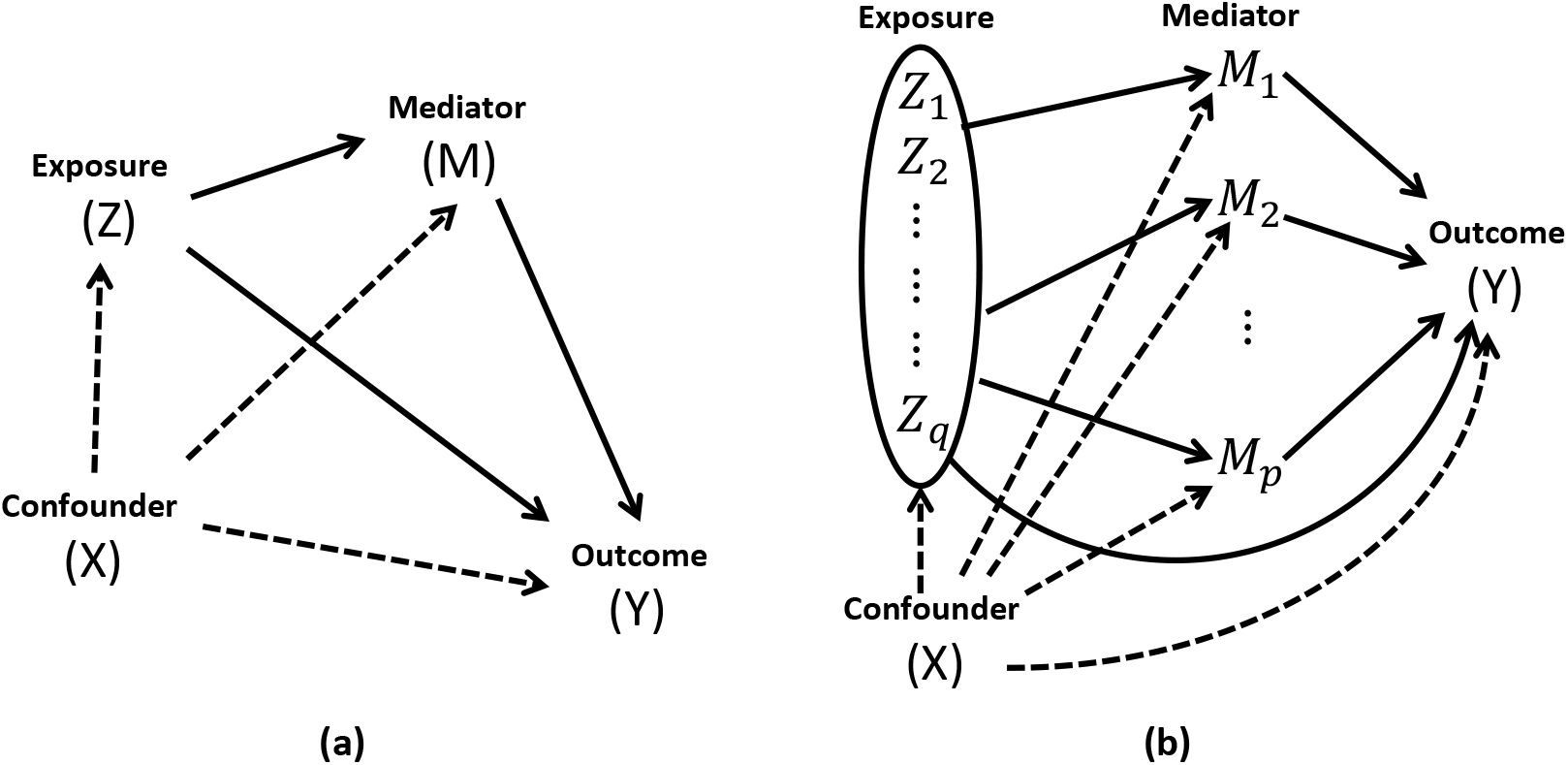
Diagram of (a) Mediation analysis with univariate exposure and univariate mediator, and (b) Mediation analysis with high dimensional exposure and high dimensional mediators considered in this paper. The solid lines are links associated with the causal effects.

The classical mediation analysis focuses on the case of univariate exposure and univariate mediator. The statistical causal inference literature has made tremendous progress in the estimation, testing and understanding of the mediation effect in such settings (Baron and Kenny, 1986; MacKinnon, 2012; VanderWeele, 2011, 2015). Mediation models with multiple mediators or multiple exposures have also been studied. For example, VanderWeele and Vansteelandt (2014) proposed two analytic approaches for multiple mediators. Huang *et al*. (2014) studied the joint analysis of the phenotype, the expression of a gene and the SNPs using mediation framework with the gene expression as the univariate mediator and the SNPs within the gene as the low-dimensional multivariate exposure. Recently, models with high dimensional mediator have began to draw attention. Huang and Pan (2016) proposed a significance test for the effect of high dimensional mediator. Chén *et al*. (2017) and Zhao *et al*. (2018) proposed PCA-like concepts for summarizing the effects of high dimensional mediators in brain imaging studies. Zhang *et al*. (2016) and Sohn *et al*. (2019) studied the sparse estimation and testing of the high dimensional mediators for univariate exposure. The focus of this paper is the sparse estimation and evaluation of high dimensional mediator effects with high dimensional exposure (Figure 1b), which is different from all above.

One motivating example is the causal gene identification problem. How the genetic variants influence the phenotypes, and what molecular mechanism mediates such effects are the central questions in genomics. Along this direction, researchers have proposed numerous methods for identifying the *causal variant* at SNP level (Kichaev *et al*., 2014) or the *causal gene* at transcriptomic level (Huang and Pan, 2016; Gamazon *et al*., 2015). We focus on the latter problem in the natural genetic context. Specifically, we are interested in finding genes directly involved in the transcriptomic pathway from genetic variants to phenotype. Mediation models have been applied to this problem, most of which modeled one gene at a time (Zhu *et al*., 2016; Barfield *et al*., 2017), or only considered univariate exposure (Huang and Pan, 2016; Zhang *et al*., 2016; Sohn *et al*., 2019).

In this paper, we study the mediation analysis with high dimensional mediator and high dimensional exposure. In particular, we are interested in mediator selection and evaluating the effect of the selected mediators. We propose two regularized regression based approaches for this high dimensional mediator selection problem, and develop relevant statistical tests for causal effects and measures of the mediation effect size. The first proposal is a new application of adaptive lasso (Zou, 2006) after modification under the conventional fixed effect regression framework for mediation analysis. The second approach is a novel method based on high dimensional linear mixed model. This proposal treats the direct effect as random effect, which reduces the model complexity and improves robustness.

The rest of the paper is organized as follows. In Section 2, we start with the model setup for the mediation analysis with high dimensional mediator and high dimensional exposure, and introduce a new measure of mediation effect adapted for high dimensional exposure. In Section 3, we proceed to our two proposed procedures, including a new variable selection algorithm for high dimensional linear mixed model. In Section 4.1, we apply the proposed methods to causal gene identification from an f2 mouse dataset for diabetes study. In Section 4.2, we evaluate the two frameworks using extensive data driven simulations. We conclude in Section 5 with discussions and future work.

## 2 Causal Mediation Model with High Dimensinal Exposure and Mediator Candidates

We are interested in the effect of a multivariate exposure vector **Z** = (*Z*_1_, …, *Z_q_*) on a univariate outcome *Y*, and how it is mediated by the candidate mediators **M** = (*M*_1_, … , *M_p_*) after being adjusted by covariates **X** (Figure 1b). In the following, we formally introduce the model and define the causal effects for mediation models with high dimensional exposure and high dimensional candidate mediators using the counterfactual framework (Rubin, 1974; VanderWeele, 2015).

Let *Y*(**z, m**) denote the outcome when the exposure vector is set to **z** and the mediator vector is set to **z**, and **M**(**z**) denote the value of the mediator vector if the exposure vector **Z** is set to **z**. The total effect (TE) when the exposure is changed from **z**\* to **z** is

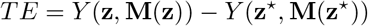

It can be decomposed in the sum of the natural indirect effect (NIE)

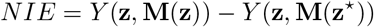

and the natural direct effect (NDE)

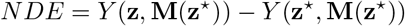

Here NIE captures the exposure effect on the outcome through the mediator, and NDE can be viewed as the controlled direct effect (CDE= *Y*(**z, m**) − *Y*(**z*, m**)) when the value of the mediator vector is set to its natural value for exposure value *z*\*. To identify these causal effects, some standard causal assumptions for mediation model are necessary (Supplementary Table 1).

In this paper, we consider the following linear models for high dimensional causal mediation.

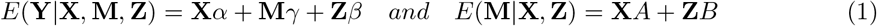

The first equation in (1) is the *outcome model* and the rest are the *mediator models*. The motivating causal gene identification problem can be formulated as mediator selection, i.e., identifying *j* ∈ {1, …, *p*} such that *γ_j_* ≠ 0 and *B_j_* ≠ 0 where *B_j_* is the *j*th column of *B*. This is the primary goal of this paper.

Another goal of this paper is to evaluate the mediation effect. When the causal assumptions and model (1) hold, the total, natural indirect and natural direct effects are reduced to

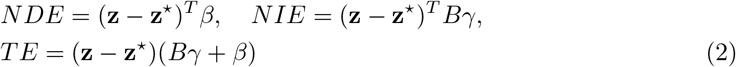

and the contribution of the individual mediators can also be defined as

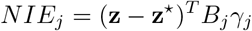

The size of the overall mediation effect is commonly defined as *PM* = *NIE/TE* in the literature (VanderWeele, 2015). For univariate exposure, *PM* is reduced to the ratio of the estimated coefficients of (1), as (*z* − *z*\*) in NIE and TE are canceled out. For multivariate exposure, if the only goal is testing whether TE, NDE or NIE exists, the test statistics do not explicitly depend on the specific realization of the exposure neither, as these tests are on the regression coefficients (Huang *et al*., 2014). In our applications, we are interested in both testing the existence of the mediation effects and estimating its size. For the latter problem, the effect size measures in (2) and PM are all functions of (*z,z*\*). They could be potentially useful in genomic selection or personalized intervention when a specific exposure combination may be of interest. However, a population level overall measure of the mediation effect size is still needed to facilitate model assessment and interpretation in scientific research practice.

We propose to define such measure by treating the exposure as a random variable. In (2), let *z*^*^ be a baseline exposure and kept fixed, and *z* be a randomly chosen exposure from the population under investigation. We use the variance of TE, NDE and NIE in (2) over the randomness of *z* as the population-level measures of effect sizes, and refer to them as the variance total effect (VTE), the variance direct effect (VDE) and the variance indirect effect (VIE). It immediately follows that

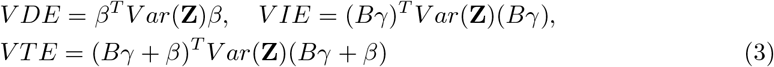

Then we define the proportion of mediation effect simply as the **P**roportion of the **V**ariance **M**ediated (PVM), i.e.,

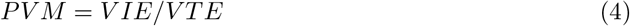

We remark that VTE is not necessarily the sum of *V IE* and *V DE* due to the potential correlatedness of the estimated direct and the indirect effects.

## 3 Regression Frameworks for High Dimensional Mediation Analysis

With some abuse of notations, let *Y* denote the observed *n* × 1 vector of the response, *Z* the *n* × *q* exposure matrix, *M* the *n* × *p* potential mediator matrix, and *X* the *n* × *s* confounder matrix including the intercept. We assume that the columns of *M* and *Z* are centered and standardized, and potentially *q, p* ≫ *n*, but presumably *s* < *n*. In genetics, *Z* could be the genotype matrix, *M* the gene expression matrix, and *X* includes the baseline covariates such as gender and age. Using the above notations, model (1) could be estimated using the following linear regression framework

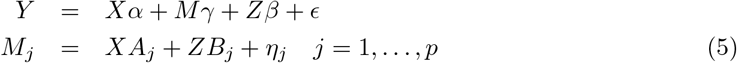

where *M_j_* is the *j*th column of matrix *M*, *ϵ* ~ *N*(0, *σ*^2^*I_n_*), and 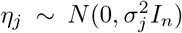. We will introduce two regression-based methods for mediation analysis with high dimensional exposure and mediators. Each procedure is consist of three steps: (a) parameter estimation by fitting the outcome model and mediator models separately, (b) causal effect testing, and (c) estimating *PVM* to measure the mediation effect.

### 3.1 MedFix: Fixed Effect Model for Mediation

#### 3.1.1 Parameter Estimation

One naive solution of (5) is applying a sparse regression technique. For the mediator models, various sparse regression procedures can be applied directly, such as adaptive lasso. For the outcome model, the predictors are heterogeneous with *M* being continuous and *Z* being discrete, and it is unclear whether such heterogeneity will cause any bias in the joint sparse selection. Thus we propose to introduce an additional tuning parameter to adjust such heterogeneity. In detail, we minimize the objective

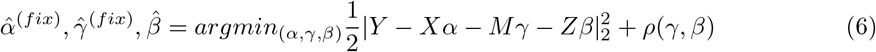

for the outcome model where

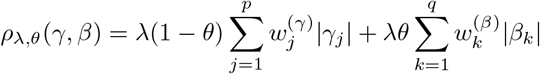

Here *λ* > 0 is the overall penalty tuning parameter, θ ∈ (0, 1) adjusts the regularization levels on the two data types, and the weight vectors *w*^(*γ*)^ and *w*^(*β*)^ are normalized separately to be summed to *p* and *q*, respectively. It can be reformulated as adaptive lasso. The mediator models can be solved by directly applying adaptive lasso. The variable weight vectors in all cases are based on the initial lasso estimates of the regression coefficients (See Supplementary Notes for details). We refer to this naive solution to (5) using existing sparse regression procedure as **Med**iation analysis via **Fix**ed effect model (**MedFix**).

#### 3.1.2 Causal Effect Testing

Next we will perform statistical tests for NIE, NDE, TE and the individual mediator effects *NIE_j_*’s. Let 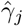 and 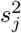 be the estimated *γ_j_* and the associated variance, and let 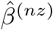 and 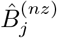 be the subvectors of the non-zero elements of the estimates of *β* and *B_j_* by MedFix, respectively, and Σ^(*β*)^ and Σ^(*B_j_*)^ their corresponding estimated variances. The tests for the causal effects can be reformulated as tests on the regression coefficients, for which we will take advantage of the oracle property of adaptive lasso (Zou, 2006). We remark that a similar testing strategy was used in Zhang *et al*. (2016) where the exposure was univariate, while we focus on high dimensional exposure in this paper.

Rejecting *H*_0, *j*_ : *NIE_j_* = 0 requires rejecting both of *H*_0, *γ_j_*_ : *γ_j_* =0 and *H*_0, *B_j_*_ : *B_j_* = 0. Let *P_γ_j__* and *P_B_j__* be their p-values, respectively. We define the p-value for *H*_0, *j*_ as *P_med,j_* = *min*(1, *P_γ_j__* + *P_B_j__*). For MedFix, *P_γ_j__* is given by a z-test as 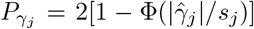 if 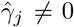, and *P_γ_j__* = 1 otherwise. *P_B_j__* is from the *χ*^2^ likelihood ratio test comparing the selected mediator model for *M_j_* and the corresponding reduced model with only *X*. Holm–Bonferroni method (Holm, 1979) is applied to *P_med,j_* for 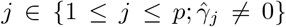 to adjust the multiplicity in mediator selection.

Testing NDE is equivalently to testing *H*_0, *β*_ : *β* = 0. We calculate its p-value from a *χ*^2^-test as 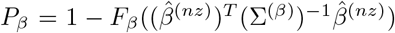 where *F_β_* is the cdf of a *χ*^2^ distribution with degree of 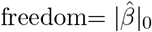.

We reject *H*_0, *med*_ : *NIE* = 0 if any of *H*_0, *j*_ is rejected. Thus its p-value is defined as *P_med_* = min(*P*_*med*,1_, …, *P_med,p_*). We reject *H*_0, *TE*_ : *TE* =0 if any of *H*_0, *med*_ and *H*_0, *β*_ is rejected, i.e., there is no total exposure effect if and only if there is neither direct effect nor indirect effect. Thus its p-value is given by *P_TE_* = min(*P_med_*, *Pβ*).

#### 3.1.3 *PVM* Estimation

Let 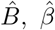 and 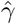 be the estimates of the coefficients in (5). Since *Var*(**Z**) can be estimated using *n*^−1^*Z^T^Z*, (3) can be estimated by 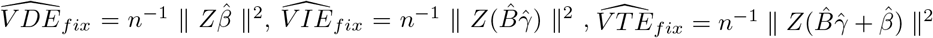 and 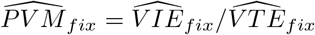.

### 3.2 MedMix: Mixed Effect Model for Mediation

In (5), *γ* models the association between the mediators and the outcome, and *β* and *B_j_*’s model the association between the exposure and the outcome/mediators, respectively. The sparse assumptions on *β* and *B_j_*’s are crucial for the initial estimation step of MedFix, but not necessary for hypothesis testing, as *H*_0, *β*_ and *H*_0, *B_j_*_ are hypotheses on the whole regression coefficient vectors instead of their individual elements. Thus an alternative strategy is to model the effects of *Z* in (5) as random effects to reduce the dimension of the parameter space in sparse selection, which lead to the following high dimensional linear mixed models

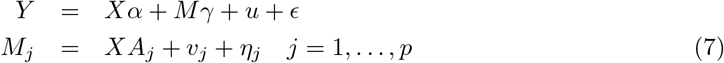

where *ϵ* ~ *N*(0, *σ*^2^*I_n_*), 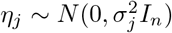 are noises, and *u* and *v_j_* for *j* = 1, …, *p* are all length *n* random genetic effect vectors such that *u* ~ *N*(0, *τK*(*Z*)) and *v_j_* ~ *N*(0, *τ_j_K*(*Z*)).

The formulation of (7) is inspired by the quantitative genetics literature where modeling the genotype-trait association as random effects has led to huge success (Hayes *et al*., 2009; Riedelsheimer *et al*., 2012). In quantitative genetics, *K*(*Z*), the covariance matrix of the random effect, is the marker-based genetic relatedness matrix of the subjects (VanRaden, 2008; Endelman and Jannink, 2012), and it could be as simple as *q*^−1^*ZZ^T^*.

#### 3.2.1 Parameter Estimation

For model (7), the mediator selection for the outcome model is related to fixed effect selection for high dimensional linear mixed model, which can be solved by minimizing the following penalized negative log-likelihood

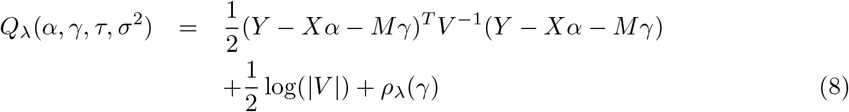

where *V* = *τK*(*Z*) + *σ*^2^*I_n_*. For this potentially non-convex problem, we propose a novel variable selection algorithm which we will discuss in Section 3.3. Together with the mediator equations in (7) fitted using conventional linear mixed model technique (e.g., R package *rrBLUP*, Endelman 2011), we term this mediation analysis procedure based on (7) as **Med**iation analysis via **Mix**ed effect model (**MedMix**).

#### 3.2.2 Causal Effect Testing

Causal effect testing for MedMix is similar to the workflow for MedFix, except the following differences. For testing the individual mediator effect, we replace *H*_0, *B_j_*_ : *B_j_* =0 with *H*_0, *τ_j_*_ : *τ_j_* = 0, for which the p-value *P_τ_j__* is calculated using a SKAT-like score test (Wang *et al*., 2011). For testing NDE, we propose to test *H*_0, *τ*_ : *τ* = 0 with its p-value *P_τ_* given by a similar score test instead of testing *H*_0, *β*_ : *β* = 0. Consequently, the p-value for *H*_0, *j*_ is *P_med,j_* = *min*(1, *P_γ_j__* + *P_τ_j__*), and the p-value for *H*_0, *TE*_ is *P_TE_* = min(*P_med_, P_τ_*).

#### 3.2.3 *PVM* Estimation

Let 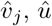 and 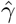 be the estimated random effects and the regression coefficients in (7). Note that in (7), *v_j_* and *u* replace *ZB_j_* and *Zβ* in (5). Define an *n* × *p* matrix 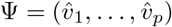, the MedMix based estimates of the causal effects in (3) are 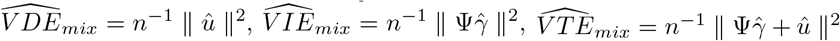 and 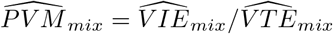.

### 3.3 Fixed Effect Selection Algorithm for MedMix

In this section, we discuss our proposed novel fixed effect selection algorithm for high dimensional linear mixed model that was deployed by MedMix for the outcome model in (7).

Fixed effects selection has been studied in the literature (Schelldorfer *et al*., 2011; Fan and Li, 2012; Müller *et al*., 2013; Rohart *et al*., 2014; Xu *et al*., 2015; Ghosh and Thoresen, 2018; Pan and Shang, 2018; Tan *et al*., 2018), most of which were on clustered data. For example, the pioneering work by Schelldorfer *et al*. (2011) proposed an *L*1 penalized estimation procedure for the fixed effect selection in high-dimensional linear mixed models. Focusing on the case of clustered data, they implemented their algorithm based on coordinate gradient descent for general random effect covariance structure, and distributed it as the R CRAN package *lmmlasso*. The model considered in this paper is different from the majority of the literature as there are no predefined clusters.

Fixed effect selection for linear mixed model naturally require the estimation of the variance components. Thus it is also related to the joint estimation of the regression coefficient and the noise level for linear model. Schelldorfer *et al*. (2011) can be regarded as such a procedure by maximizing the joint likelihood of all parameters. For *iid* noise, Sun and Zhang (2010, suggested that such one-stage maximum likelihood approach may cause bias in noise level estimation, and the proposed scaled lasso that iterates between estimating the regression coefficients and the noise level. Their procedure enjoys joint convexity, and the solution is almost always unique (Section 2 of Sun and Zhang 2012). Motivated by their success, we adopt a similar alternating optimization strategy (Bezdek and Hathaway, 2003) with a scaled adaptive lasso penalty

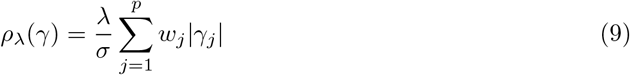

where *w_j_*’s are the weights of the variables scaled to be summed to *p*.

Consider the objective function (8), with *V* = *τK*(*Z*) + *σ*^2^*I_n_* and the scaled adaptive lasso penalty (9). Suppose *K*(*Z*) is full-rank, and let its eigen decomposition be *UDU^T^*. It follows that *V* = *UD*(*τ, σ*^2^)*U^T^* where *D*(*τ, σ*^2^) = *diag*(*τd*_1_ + *σ*^2^, …, *τd_n_* + *σ*^2^) and *d_i_*’s are the known eigen values of *K*(*Z*). Let *Ỹ* = *U^T^Y*, 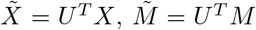, and *ỹ_i_*, 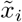, and 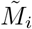 be their *i*’th row, respectively. (8) can be re-written as

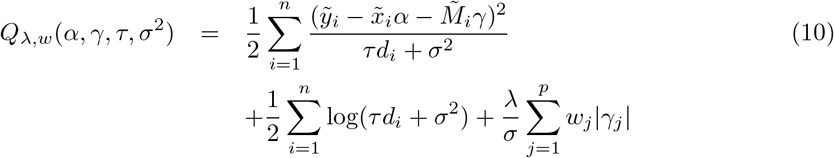

Reparametrize the above loss function by defining

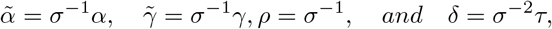

then (10) becomes

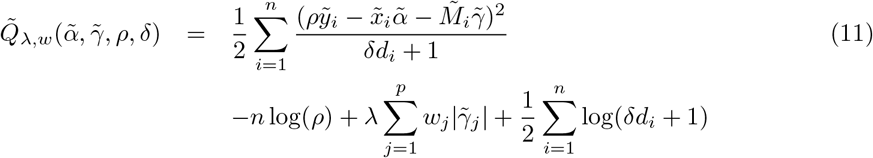

It is jointly convex when *d_i_*’s are all 0, corresponding to the case with iid noise as for the original scaled lasso. However, when the noise are correlated as in the linear mixed model under consideration, the joint convexity of this objective function depends on the parameters, the data and *d_i_*’s in a complicated way. Thus the direct optimization of (11) is not guaranteed to reach a local minimum, even though using a coordinate gradient descent assures the convergence to a stationary point (Schelldorfer *et al*., 2011).

In fact, (11) is the sum of a concave function (the last term) and a convex function (the sum of the others), for which a concave-convex procedure (CCCP, Yuille and Rangarajan 2003) can be applied. Consider an objective *Q*(*x*) = *Q_vex_*(*x*) + *Q_cav_*(*x*) where *Q_vex_*(*x*) and *Q_cav_*(*x*) are a convex and a concave function, respectively. Let 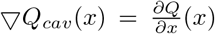, and *x*^(*t*)^ be the current estimate of *x*, then *Q*^(*t*)^(*x*) = *Q_vex_*(*x*) + *x* · ▿*Q_cav_*(*x*^(*t*)^) is a convex tight upper bound of *Q*(*x*). CCCP minimizes this upper bound in each iteration, and eventually converges to a local minimum of the original objective. It has been used in the literature for implementing SCAD (Fan and Li, 2001; Kim *et al*., 2008).

For our algorithm, let 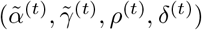 be a current estimate of 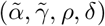, a tight upper bound of (11) is

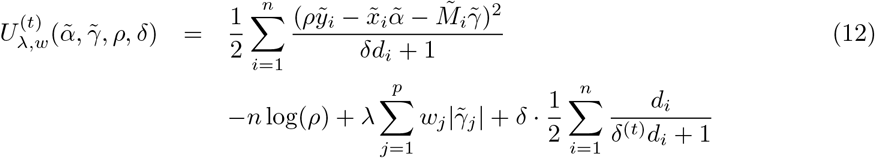

In each iteration, we minimize this upper bound by alternatively updates the regression coefficients and the variance components related parameters.

## 4 Results

MedFix has two tuning parameters (*λ, θ*). When *θ* = 0.5, the penalty on the two data types in the outcome model of (5) are the same, and *λ* can be determined by minimizing *BIC*. We call this version of MedFix MedFix_0.5_. Alternatively, (*λ, θ*) can be jointly selected by BIC, which we term as MedFix_*BIC*_. MedMix has only one tuning parameter *λ*, and we determine it using *BIC* (Schelldorfer *et al*., 2011). The random effect covariance matrix *K*(*Z*) can be modeled in many ways. Since our application is in genetics, we deploy two estimates of the genetic relatedness matrix, the original proposal (VanRaden, 2008) and a shrinkage estimate (Endelman and Jannink, 2012). MedMix with these two types of covariance matrix for the random effect are referred to as MedMix_*linear*_ and MedMix_*shrink*_, respectively. We will compare these four protocols in real data analysis and data-driven simulations. Throughout this paper, we set the significance level for mediator selection at 0.1.

### 4.1 Evaluation based on Real Data

We analyze a mouse f2 cross data for diabetes study (Wang *et al*., 2011; Tu *et al*., 2012; Tian *et al*., 2015). This dataset include the genotype captured by SNP array with 2057 markers, microarray-based gene expression with about 40k probes from six tissues, and various clinical phenotypes. The phenotypes we used as outcome are the plasma insulin level at 10 weeks before sacrifice. We are interested in identifying potential causal genes for insulin level. Since insulin is secreted from islet, we use the islet gene expression as the candidate mediator, and genotype as the exposure. We use gender as a covariate in all mediator models and outcome models. There are 491 mice with all three data types. One unique feature of this dataset is that it includes the following three independent measures of plasma insulin at 10 week which we will refer to as **IA**, **IB**, and **IC**:

**IA** : “INSULIN (ng/ml) 10 wk” from lipomics measure, the primary insulin measurement.
**IB** : “insulin 10 wk” from lipomics, a reproduced measurement
**IC** : “Insulin (uIU/mL)”from RBM panel, a measurement using a different technology.

This enables us to compare methods based on the reproducibility of the analysis results across these phenotypes representing the independent measures of biologically identical signal, in the absence of “grand truth” in real data. We do not filter the genotype markers, but do screen the microarray probes, and only keep those share QTL and have reasonable correlations (|*corr*| ≥ 0.05) with the outcome (See Supplementary Notes for details of data preprocessing). We primarily focus on the accuracy and reproducibility in mediator selection and PVM estimation in the main paper, and the testing results on NIE, NDE and TE are in Supplementary Table 2.

#### 4.1.1 MedMix tend to select more relevant mediators

Since there is no well-established “grand truth” of causal genes for insulin, we use the known biological annotations as a proxy for evaluating the causal gene identification accuracy. For each gene, we record whether or not it is involved in a Gene Ontology (GO) term (Consortium, 2004), KEGG pathway (Kanehisa and Goto, 2000) or Medical Subject Headings (MeSH) term (Lipscomb, 2000) for insulin, diabetes or islet/pancreas function. In Table 1, we present the numbers of the mediators selected for each insulin measure by each method, and how many of them belong to a relevant term. We remark that this is not an enrichment analysis, and it is only based on their presence or absence in relevant terms. We find that MedMix_*shrink*_ has the highest proportion of selected mediators with such annotations for phenotypes IA and IC, MedFix_0.5_ is the highest for IB.

**Table 1:**
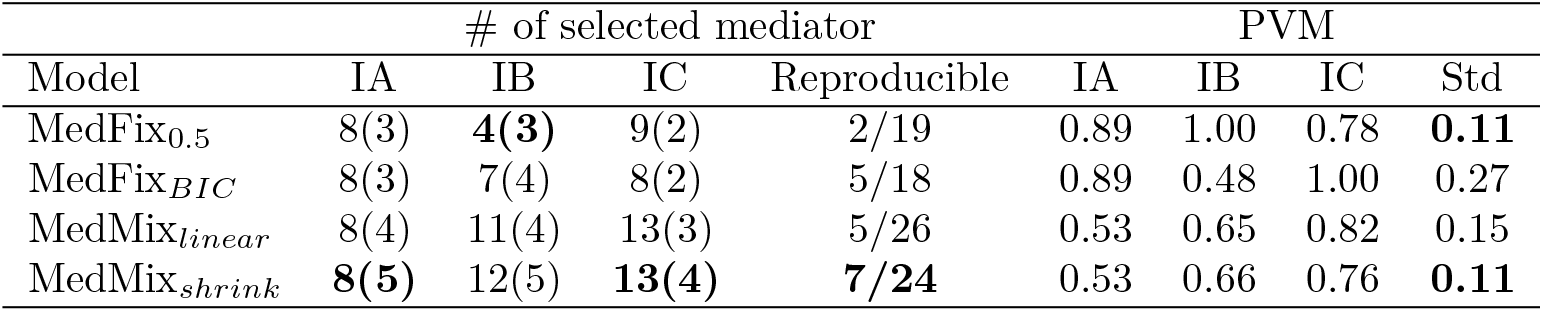
Analysis of three independent measurements of insulin level before sacrifice. For the columns of # of selected mediator genes, the numbers in parenthesis are the number of genes known to be relevant to pancreas function and/or diabetes determined by their affilication to the relevant GO/KEGG/MeSH terms. In each column of IA, IB and IC, the method with the highest proportion of relevant genes is in boldface. The column Reproducible presents the number of mediators selected by at least two of the three phenotypes, and the total number of genes selected for any phenotype. The method with the highest such ratio is boldfaced. We present in the column Std the standard deviation of the estimated PVM’s across three phenotypes for each method, and boldface the smallest ones.

#### 4.1.2 MedMix yield more reproducible mediators and estimates of overall mediation effect size

Since the three phenotypes measure the same biological signal, a good mediator selection procedure should output similar genes in these analyses. We refer to a mediator as “reproducible” if it is selected in at least two of the three analysis, and find that MedMix_*shrink*_ performs the best in terms of the ratio of the reproducible mediators and the mediator selected in any analysis, and MedFix_*BIC*_ scores the second (Table 1). We also calculate the variance of the estimated PVM’s across the three phenotypes for each method, and find that MedFix_*BIC*_ leads to very large variation, while the other three methods are comparable. Overall, we conclude that MedMix_*shrink*_ yields the most reproducible results.

#### 4.1.3 MedMix and MedFix can differentiate causal mediation effect and pleiotropy

It is known that insulin is secreted by islet. Thus the islet gene expressions are its most direct transcriptomic mediators. If the gene expressions from another tissue are used as the candidate mediator, the causal assumptions for mediation analysis would be violated. Any mediator genes selected in such analysis is most likely due to pleiotropy, spurious correlation or causal effect with reverse direction. It is difficult, if not impossible, for any quantitative model to automatically investigate the scientific appropriateness of the mediator candidates. Nevertheless, it is reasonable to expect a much smaller estimated PVM when irrelevant mediator candidates are used. We run MedFix and MedMix using gene expressions from each of the other five tissues (adipose, gastroc, hypothalamus, kidney and liver) as the candidate mediators, and calculate the corresponding PVM. If the output PVM from an irrelevant tissue is equal to or larger than the corresponding islet PVM, we label this case as a “failure” of separating the true and spurious causal mediation effects. We find that all methods output smaller PVMs in most cases when an irrelevant tissue is used as the potential mediators (Table 2). In particular, MedFix_0.5_ and MedMix_*shrink*_ fail less often than the other two. Most failures are for adipose and gastroc. It makes biological sense, because plasma insulin directly and indirectly (through glucose) regulates the expressions of a wide range of genes in adipose and gastroc (Ducluzeau *et al*., 2001; Elbein *et al*., 2011). It reminds us that mediation analysis cannot distinguish the true causal mediation and the causal effect with reverse direction, which is a fundamental limitation of mediation models of all kinds.

**Table 2:** Estimated PVM using gene expressions from various tissues. Only islet gene expressions are expected to directly regulate insulin. The cases where the reported PVM is larger than the PVM from the islet case are boldfaced.

| Outcome IA | Islet | Adipose | Gastroc | Hypothalamus | Kidney | Liver |
| --- | --- | --- | --- | --- | --- | --- |
| MedFix<sub>0.5</sub> | 0.89 | **1.00** | 0.57 | 0.50 | 0.55 | 0.32 |
| MedFix<sub>BIC</sub> | 0.89 | 0.63 | **1.00** | 0.65 | 0.24 | 0.26 |
| MedMix<sub>linear</sub> | 0.53 | **0.70** | 0.32 | 0.19 | 0.30 | 0.24 |
| MedMix<sub>shrink</sub> | 0.53 | **0.89** | 0.33 | 0.22 | 0.33 | 0.29 |
| Outcome IB | Islet | Adipose | Gastroc | Hypothalamus | Kidney | Liver |
| MedFix<sub>0.5</sub> | 1.00 | **1.00** | 0.54 | 0.28 | 0.25 | 0.62 |
| MedFix<sub>BIC</sub> | 0.48 | **0.62** | **0.78** | 0.10 | 0.25 | **1.00** |
| MedMix<sub>linear</sub> | 0.65 | **0.75** | 0.38 | 0.14 | 0.32 | 0.21 |
| MedMix<sub>shrink</sub> | 0.66 | **0.66** | 0.46 | 0.23 | 0.32 | 0.18 |
| Outcome IC | Islet | Adipose | Gastroc | Hypothalamus | Kidney | Liver |
| MedFix<sub>0.5</sub> | 0.78 | 0.55 | 0.37 | 0.15 | 0.36 | 0.39 |
| MedFix<sub>BIC</sub> | 1.00 | 0.22 | 0.12 | 0.15 | 0.27 | 0.51 |
| MedMix<sub>linear</sub> | 0.82 | 0.54 | 0.41 | 0.06 | 0.26 | 0.36 |
| MedMix<sub>shrink</sub> | 0.76 | 0.55 | **1.00** | 0.07 | 0.26 | 0.25 |

### 4.2 Evaluation based on Simulations

We simulate data using a pipeline based on the above f2 mouse data analysis with IA as the outcome. The goal of our simulations is investigating the accuracy of mediator selection of MedFix and MedMix, and their robustness against model mis-specification.

#### 4.2.1 Real Data Driven Simulation Model

We design a data-driven simulation model using the genotype, preprocessed islet gene expression and gender from the f2 mouse dataset as basis. We simulate the outcome *Y* and the mediators 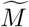 using a hybrid of (5) and (7) controlled by a simulation parameter *ϕ* ∈ [0, 1]. When *ϕ* = 0, 1, (5) and (7) are the true models, respectively. When *ϕ* ∈ (0, 1), both MedMix and MedFix are misspecified. Thus this simulation model provides a common platform for investigating their robustness against model misspecification. The other two simulation parameters that we investigate are (*h, g*), the strength of the mediation effect and the direct effect, respectively. In our simulations, we assume there are 15 true mediators, and 15 fake mediators with mediator-outcome link but without exposure-mediator link. The other ≈11k candidate mediators have no mediator-outcome link, regardless whether it is controlled by the exposure (See Supplementary Notes for details of the data-driven simulation model). We consider all 12 combinations of *g* =1, 2, *h* =1, 4 and *ϕ* = 0, 0.5, 1. For each scenario, we repeat the simulation for 40 replicates.

#### 4.2.2 MedMix is more accurate and robust in mediator selection

We compare the mediator selection accuracy using the simulated data (Table 3), and find that in most of cases, MedMix models yield less false positives than MedFix methods with a slight increase in false negatives. The overall mediator selection errors (the sum of false positives and false negatives) for MedMix are almost always smaller than those for MedFix. In detail, when *ϕ* = 0, the simulation model is the same as the model underlying MedFix (5), and MedMix is mis-specified. However, MedFix and MedMix have comparable overall mediator selection errors. In contrast, when *ϕ* =1 and the MedMix model (7) is the true model, MedMix clearly lead to less errors in all settings. When *ϕ* = 0.5, neither (5) nor (7) is the true model, and both of MedFix and MedMix are misspecified. In these cases, MedMix also perform better than MedFix in all scenarios. We further compare their variable selection accuracy for the outcome models (identifying nonzero elements of *γ*) without any significance tests, and observe similar advantage of MedMix (Supplementary Table 3). These results suggest that MedMix yields more accurate results than MedFix in a wider range of settings, and is more robust against model misspecification.

**Table 3:** The number of of false positives and false negatives in mediator selection in simulations. There are 15 true mediators in all simulation settings.

| $(h, g, \phi)$ | ( 1 , 1 , 0 ) | ( 1 , 2 , 0 ) | ( 4 , 1 , 0 ) | ( 4 , 2 , 0 ) |
| --- | --- | --- | --- | --- |
| MedFix<sub>0.5</sub> | 1 , 0 | 1.6 , 0.1 | 2.4 , 0.1 | 1.4 , 0 |
| MedFix<sub>BIC</sub> | 1.1 , 0 | 1.5 , 0.1 | 1.7 , 0 | 1.6 , 0 |
| MedMix<sub>linear</sub> | 0.5 , 1.2 | 0.5 , 1.4 | 0.3 , 1.1 | 0 , 1.1 |
| MedMix<sub>shrink</sub> | 0.4 , 1.2 | 0.4 , 1.7 | 0.5 , 1.1 | 0 , 1.1 |
| $(h, g, \phi)$ | ( 1 , 1 , 0.5 ) | ( 1 , 2 , 0.5 ) | ( 4 , 1 , 0.5 ) | ( 4 , 2 , 0.5 ) |
| MedFix<sub>0.5</sub> | 3.1 , 0.4 | 4.1 , 2 | 2.5 , 0.1 | 4.1 , 0.2 |
| MedFix<sub>BIC</sub> | 3.5 , 0.3 | 2.8 , 1.8 | 2.9 , 0.1 | 3.7 , 0.2 |
| MedMix<sub>linear</sub> | 0.3 , 1.5 | 0.6 , 1.8 | 0.6 , 1.2 | 0.1 , 1.6 |
| MedMix<sub>shrink</sub> | 0.4 , 1.5 | 0.3 , 1.8 | 0.5 , 1.2 | 0.1 , 1.6 |
| $(h, g, \phi)$ | ( 1 , 1 , 1 ) | ( 1 , 2 , 1 ) | ( 4 , 1 , 1 ) | ( 4 , 2 , 1 ) |
| MedFix<sub>0.5</sub> | 3.2 , 1.4 | 3 , 6.1 | 3.9 , 0.2 | 3.9 , 0.4 |
| MedFix<sub>BIC</sub> | 3.3 , 1.3 | 3.2 , 6 | 2.9 , 0.2 | 3.7 , 0.4 |
| MedMix<sub>linear</sub> | 0.6 , 1.5 | 0.4 , 2.9 | 0.2 , 1.4 | 0.1 , 1.5 |
| MedMix<sub>shrink</sub> | 0.4 , 1.5 | 0.4 , 2.8 | 0.3 , 1.4 | 0.1 , 1.5 |

## 5 Discussion

In this paper, we study the mediation analysis with high dimensional candidate mediator and high dimensional exposure, and its application to causal gene detection. We develop two procedures for mediator selection and their associated causal effect tests and mediation effect measure. MedFix is an application of adaptive lasso with an additional tuning parameter for the outcome model balancing the penalties on the exposure and the mediator variables. MedMix is based on high dimensional linear mixed models, for which we develop a new fixed effect selection algorithm for the outcome model. In our data driven simulation studies, we show that MedMix lead to more accurate mediator selection, and are more robust against model misspecification. We apply these methods to causal gene identification from an f2 mouse dataset for diabetes study, and find that MedMix is more likely to select higher proportions of relevant genes, and yields more reproducible results in mediator selection and PVM estimation.

We focus on mediator selection and the evaluation of the joint mediation effect of the selected mediators in this paper, and do not discuss the roles of the individual mediators. Due to the potential mediator-mediator interaction, quantifying such individual effects requires stronger assumptions, even when there are only a few mediators, and their order in the causal chain is known (Daniel *et al*., 2015). Thus we leave it for future research. We do not consider the exposure-exposure interaction, exposure-mediator interaction or nonlinear causal effects neither. Generalizations along this direction could be made potentially by extending (5) to additive models or by modeling *K*(*Z*) in (7) with an appropriate kernel function (Morota and Gianola, 2014). Another future work we will pursue is to extend the current framework to the joint analysis of multiple outcomes. In the context of causal gene identification, such high dimensional mediation framework for multiple phenotype will dissect the pleiotropy of the phenotyeps, and lead to refined phenotypical causal networks with genetic basis.

## Acknowledgements

We thank Dr. Alan Attie for providing easy access to the data and insightful discussions on the biology of diabetes, and the Holland Computing Center (HCC) at UNL for computation resources and technical supports. This work has been supported by NSF ABI (Award# DBI-1564621), NSF EPSCoR (RII) Track II (Award# OIA-1736192) and NU Collaborative System Science Seed Grant to QZ.

## Supplementary Materials

### Supplementary Notes

#### Variable Weights for Adaptive Lasso

We define the weights for adaptive lasso for the mediator models of MedFix as the following.

For mediator *M_j_*, first apply regular lasso with tuning parameter *λ*, and no penalty on *λ_j_* to estimate (*λ_j_*, *B_j_*) in the mediator equation

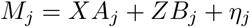

let 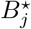 be this initial estimate. Then we define the weights 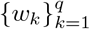 for the final adaptive lasso estimation with the parameters (*λ, θ*) such that 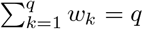 and

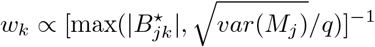

where 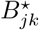 is the *k*th element of 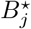. The intuition behind this weight is that if the initial estimate of a nonzero coefficient is small enough 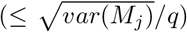, it will have the same weight as the zero elements.

Similarly, we define the weights for the outcome models of MedFix as the following. Let 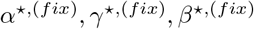 be the solution of (6) with penalty

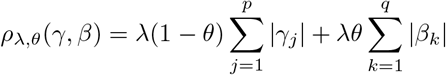

for fixed *λ* and *θ*.

Then we define the weights 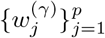 and 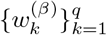 for the final adaptive lasso estimation with parameter (*λ, θ*) such that 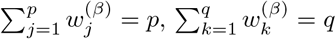,

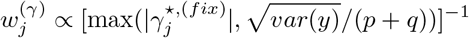

and

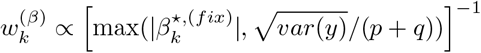

For the outcome model of MedMix, the weights are updated in each iteration as the following. Let

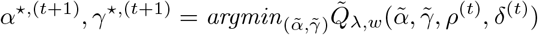

where all elements of the length *p* vector *w* are all 1. We define variable weights 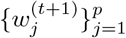 such that 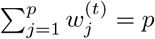 and

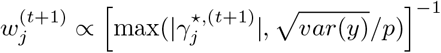

#### F2 mouse data preprocessing

Mediation effect implies the existence of the association between the response and the exposure, the mediator and the exposure, and the mediator and the response. We use these assumptions to filter genes that enter our analysis. We only keep genes that satisfies the following three criteria. (1) Islet expression shares QTL with the phenotype under investigation (lod score ≥ 3). (2) Islet expression correlates with the phenotype (|*corr*| ≥ 0.05). (3) Islet expression has larger correlation with the phenotype than the expression in the other five tissues. We design this filtering procedure using low thresholds with the hope of including as many genes as possible, and about 11k genes survive for each insulin measure. The QTL/eQTL information are from Dr. Attie Lab Diabetes Database (http://diabetes.wisc.edu).

#### Real Data Driven Simulation Model

We simulate the outcome *Y* and the mediators 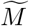 using a hybrid of MedFix model (5) and the MedMix model (7), controlled by a simulation parameter *ϕ* ∈ [0, 1]. When *ϕ* = 0, 1, MedFix and MedMix are the true models, respectively. When *ϕ* ∈ (0, 1), both MedMix and MedFix are misspecified. Thus our simulation setting provides a common platform for investigating their robustness against model misspecification. The other two simulation parameters that we investigate are (*h, g*), the strength of the mediation effect and the direct effect, respectively.

In the following, we elaborate on our data-driven simulation model. In the f2 mouse data [Wang et al., 2011, Tu et al., 2012, Tian et al., 2015] analysis of the phenotype IA as in this paper, 15 genes were selected with in the outcome model by all four methods under consideration. We will use them as the true mediators for the simulation model, and let *S_T_* ⊂ {1, …, *p*} to denote their indices. Let *S_F_* be another 15 indices evenly spaced between 1 to p which we will use as the set of fake mediators with non-zero regression coefficients in the outcome model, but no exposure-mediator link.

The following elements are fixed for simulations.

- *Z*: the 491 × 2057 normalized genotype matrix from the real data.
- *X*: the intercept and sex from the real data
- Let 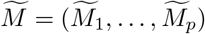 denote the mediator matrix for our simulation student. We set 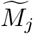 be the same as *M_j_*, the islet gene expression in the real data, for *j* ∈ *S_T_* ∪ *S_F_*, i.e., the irrelevant genes.
- *α* = (1, 0.2)^*T*^
- *β*_0_: a length 2057 sparse vector who has eight nonzero coordinates evenly spaced from 1 to 2057 with values (1,-1, 2,-2, 1,-1, 2,-2).
- *γ*_0_: a length p sparse vector, whose nonzero elements are in *S_T_* ∪ *S_F_* with values (1,-1, 2,-2, 1,-1, 2,-2, 1,-1, 2,-2, 1,-1, 2,-2, 1,-1, 2,-2, 1,-1, 2,-2, 1,-1, 2,-2, 1,-1).
- 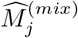 and 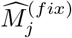, the predicted values of *M_j_* in the real data analysis based on MedMix and MedFix, respectively, for *j* ∈ *S_T_*,i.e,.

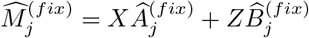

where 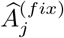 and 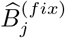 are the estimated regression coefficients by MedFix, and

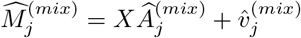

where 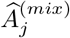 is the estimated regression coefficient and 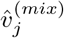 is the predicted random effect by MedMix.

For *j* ∈ *S_T_*, we simulate

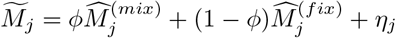

where 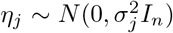 and 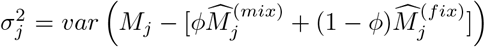. Note that in the real data analysis, the MedMix and MedFix tests on exposure-mediator link for these 15 genes all return small p-values. So it is also expected for the simulated true mediators.

For the fake mediators with no exposure-mediator link *j* ∈ *S_F_*, we first generate

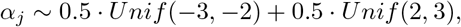

then simulate

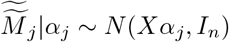

and finally normalize 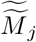 to 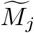 so that it has the same mean and variance as *M_j_*, the expression of gene *j* in real data.

We calculate the direct effect based on MedFix *β* = *g* · *β*_0_, and simulate the direct effect based on MedMix *u* ~ *N*(0, *τ_u_ZZ^T^*) where 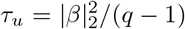. This is to assure that *u* and *Zβ*, the direct effects based on MedMix and MedFix, respectively, to have similar effect size on the outcome.

Finally, let the mediator regression coefficient *γ* = *h* · *γ*_0_, and we simulate the outcome as

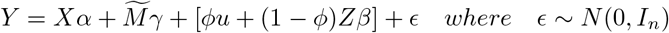

To summarize, there are three simulation parameters (*h, g, ϕ*), representing the strengths of the mediation effect, direct effect and the mixing weight of the underlying model of MedMix in above simulation model. In particular, the use of mixing weight *ϕ* in the simulation model enable us to study the robustness of MedMix and MedFix against model misspecification. We consider all combinations of *g* =1, 2, *h* = 1, 4 and *ϕ* = 0, 0.5, 1. For each scenario, we repeat the simulation for 40 replicates.

#### Supplementary Figures and Tables

**Table S1:**
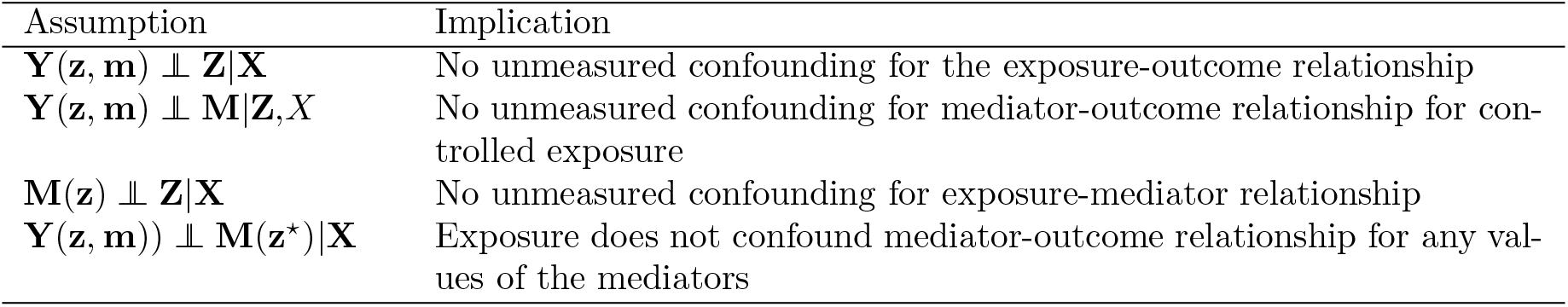
Standard causal assumptions for mediation analysis and their implications.

**Table S2:** p-values for the causal effects in the real data analysis.

| IA |  |  |  |
| --- | --- | --- | --- |
| Model | NDE | NIE | TE |
| MedFix<sub>0.5</sub> | $5.9 \times 10^{-4}$ | $1.7 \times 10^{-4}$ | $1.7 \times 10^{-4}$ |
| MedFix<sub>BIC</sub> | $1.6 \times 10^{-3}$ | $1.1 \times 10^{-4}$ | $1.1 \times 10^{-4}$ |
| MedMix<sub>linear</sub> | $3.3 \times 10^{-6}$ | $2.0 \times 10^{-5}$ | $3.3 \times 10^{-6}$ |
| MedMix<sub>shrink</sub> | $2.3 \times 10^{-5}$ | $9.9 \times 10^{-10}$ | $9.9 \times 10^{-10}$ |
| IB |  |  |  |
| Model | NDE | NIE | TE |
| MedFix<sub>0.5</sub> | 1.0 | $9.8 \times 10^{-8}$ | $9.8 \times 10^{-8}$ |
| MedFix<sub>BIC</sub> | $2.1 \times 10^{-15}$ | $4.5 \times 10^{-7}$ | $2.1 \times 10^{-15}$ |
| MedMix<sub>linear</sub> | $2.3 \times 10^{-4}$ | $1.5 \times 10^{-8}$ | $1.5 \times 10^{-8}$ |
| MedMix<sub>shrink</sub> | $1.90 \times 10^{-4}$ | $1.5 \times 10^{-7}$ | $1.5 \times 10^{-7}$ |
| IC |  |  |  |
| Model | NDE | NIE | TE |
| MedFix<sub>0.5</sub> | $5.8 \times 10^{-2}$ | $1.9 \times 10^{-5}$ | $1.9 \times 10^{-5}$ |
| MedFix<sub>BIC</sub> | 1.0 | $4.7 \times 10^{-7}$ | $4.7 \times 10^{-7}$ |
| MedMix<sub>linear</sub> | $1.9 \times 10^{-5}$ | $1.2 \times 10^{-13}$ | $1.2 \times 10^{-13}$ |
| MedMix<sub>shrink</sub> | $2.2 \times 10^{-6}$ | $7.4 \times 10^{-6}$ | $2.2 \times 10^{-6}$ |

**Table S3:** The number of of false positives and false negatives in variable selection for the outcome models in simulations. There are 30 candidate mediators with non-zero coefficient in the outcome models in all simulation settings.

| $(h, g, \phi)$ | ( 1 , 1 , 0 ) | ( 1 , 2 , 0 ) | ( 4 , 1 , 0 ) | ( 4 , 2 , 0 ) |
| --- | --- | --- | --- | --- |
| MedFix<sub>0.5</sub> | 17.4 , 1.6 | 13.2 , 5 | 11.3 , 0.1 | 11.4 , 0.3 |
| MedFix<sub>BIC</sub> | 20.2 , 1.4 | 17 , 4.7 | 8.4 , 0.1 | 11.2 , 0.3 |
| MedMix<sub>linear</sub> | 36.4 , 0.8 | 39.2 , 3.6 | 9.6 , 0.1 | 13.3 , 0.2 |
| MedMix<sub>shrink</sub> | 31.7 , 1 | 30.2 , 4.8 | 10.7 , 0.1 | 13.1 , 0.2 |
| $(h, g, \phi)$ | ( 1 , 1 , 0.5 ) | ( 1 , 2 , 0.5 ) | ( 4 , 1 , 0.5 ) | ( 4 , 2 , 0.5 ) |
| MedFix<sub>0.5</sub> | 33.6 , 2.6 | 34.5 , 7.3 | 13.1 , 0 | 19.4 , 0.4 |
| MedFix<sub>BIC</sub> | 29.7 , 2.4 | 24.4 , 7.3 | 11.3 , 0 | 16.4 , 0.4 |
| MedMix<sub>linear</sub> | 31.6 , 0.5 | 40.7 , 2.6 | 10.8 , 0.3 | 12.4 , 0.1 |
| MedMix<sub>shrink</sub> | 29.3 , 0.7 | 33.5 , 3.2 | 10.5 , 0.2 | 12.8 , 0.1 |
| $(h, g, \phi)$ | ( 1 , 1 , 1 ) | ( 1 , 2 , 1 ) | ( 4 , 1 , 1 ) | ( 4 , 2 , 1 ) |
| MedFix<sub>0.5</sub> | 37.9 , 5.9 | 31.4 , 14.7 | 19.1 , 0.2 | 28.3 , 1.6 |
| MedFix<sub>BIC</sub> | 29.7 , 5.9 | 25.6 , 15.2 | 14.3 , 0.1 | 25.2 , 1.4 |
| MedMix<sub>linear</sub> | 40.2 , 1.1 | 37.7 , 5.5 | 10.2 , 0.1 | 15.8 , 0 |
| MedMix<sub>shrink</sub> | 38.4 , 1.1 | 31.9 , 5.7 | 10.6 , 0.1 | 16.1 , 0.2 |

## References

Barfield, R., Shen, J., Just, A. C., Vokonas, P. S., Schwartz, J., Baccarelli, A. A., VanderWeele, T. J., and Lin, X. (2017). Testing for the indirect effect under the null for genome-wide mediation analyses. Genetic epidemiology, 41(8), 824–833.

Baron, R. M. and Kenny, D. A. (1986). The moderator–mediator variable distinction in social psychological research: Conceptual, strategic, and statistical considerations. Journal of personality and social psychology, 51(6), 1173.

Bezdek, J. C. and Hathaway, R. J. (2003). Convergence of alternating optimization. Neural, Parallel & Scientific Computations, 11(4), 351–368.

Chén, O. Y., Crainiceanu, C., Ogburn, E. L., Caffo, B. S., Wager, T. D., and Lindquist, M. A. (2017). High-dimensional multivariate mediation with application to neuroimaging data. Biostatistics, 19(2), 121–136.

Consortium, G. O. (2004). The gene ontology (go) database and informatics resource. Nucleic acids research, 32(suppl_1), D258–D261.

Daniel, R., De Stavola, B., Cousens, S., and Vansteelandt, S. (2015). Causal mediation analysis with multiple mediators. Biometrics, 71(1), 1–14.

Ducluzeau, P.-H., Perretti, N., Laville, M., Andreelli, F., Vega, N., Riou, J.-P., and Vidal, H. (2001). Regulation by insulin of gene expression in human skeletal muscle and adipose tissue: evidence for specific defects in type 2 diabetes. Diabetes, 50(5), 1134–1142.

Elbein, S. C., Kern, P. A., Rasouli, N., Yao-Borengasser, A., Sharma, N. K., and Das, S. K. (2011). Global gene expression profiles of subcutaneous adipose and muscle from glucose-tolerant, insulin-sensitive, and insulin-resistant individuals matched for bmi. Diabetes, page DB_101270.

Endelman, J. B. (2011). Ridge regression and other kernels for genomic selection with r package rrblup. The Plant Genome, 4(3), 250–255.

Endelman, J. B. and Jannink, J.-L. (2012). Shrinkage estimation of the realized relationship matrix. G3: Genes, Genomes, Genetics, 2(11), 1405–1413.

Fan, J. and Li, R. (2001). Variable selection via nonconcave penalized likelihood and its oracle properties. Journal of the American statistical Association, 96(456), 1348–1360.

Fan, Y. and Li, R. (2012). Variable selection in linear mixed effects models. Annals of statistics, 40(4), 2043.

Gamazon, E. R., Wheeler, H. E., Shah, K. P., Mozaffari, S. V., Aquino-Michaels, K., Carroll, R. J., Eyler, A. E., Denny, J. C., Nicolae, D. L., Cox, N. J., et al. (2015). A gene-based association method for mapping traits using reference transcriptome data. Nature genetics, 47(9), 1091.

Ghosh, A. and Thoresen, M. (2018). Non-concave penalization in linear mixed-effect models and regularized selection of fixed effects. AStA Advances in Statistical Analysis, 102(2), 179–210.

Hayes, B. J., Bowman, P. J., Chamberlain, A., and Goddard, M. (2009). Invited review: Genomic selection in dairy cattle: Progress and challenges. Journal of dairy science, 92(2), 433–443.

Holm, S. (1979). A simple sequentially rejective multiple test procedure. Scandinavian journal of statistics, pages 65–70.

Huang, Y.-T. and Pan, W.-C. (2016). Hypothesis test of mediation effect in causal mediation model with high-dimensional continuous mediators. Biometrics, 72(2), 402–413.

Huang, Y.-T., VanderWeele, T. J., and Lin, X. (2014). Joint analysis of snp and gene expression data in genetic association studies of complex diseases. The annals of applied statistics, 8(1), 352.

Kanehisa, M. and Goto, S. (2000). Kegg: kyoto encyclopedia of genes and genomes. Nucleic acids research, 28(1), 27–30.

Kichaev, G., Yang, W.-Y., Lindstrom, S., Hormozdiari, F., Eskin, E., Price, A. L., Kraft, P., and Pasaniuc, B. (2014). Integrating functional data to prioritize causal variants in statistical fine-mapping studies. PLoS genetics, 10(10), e1004722.

Kim, Y., Choi, H., and Oh, H.-S. (2008). Smoothly clipped absolute deviation on high dimensions. Journal of the American Statistical Association, 103(484), 1665–1673.

Lipscomb, C. E. (2000). Medical subject headings (mesh). Bulletin of the Medical Library Association, 88(3), 265.

MacKinnon, D. (2012). Introduction to statistical mediation analysis. Routledge.

Morota, G. and Gianola, D. (2014). Kernel-based whole-genome prediction of complex traits: a review. Frontiers in genetics, 5, 363.

Müller, S., Scealy, J. L., Welsh, A. H., et al. (2013). Model selection in linear mixed models. Statistical Science, 28(2), 135–167.

Pan, J. and Shang, J. (2018). Adaptive lasso for linear mixed model selection via profile log-likelihood. Communications in Statistics-Theory and Methods, 47(8), 1882–1900.

Riedelsheimer, C., Czedik-Eysenberg, A., Grieder, C., Lisec, J., Technow, F., Sulpice, R., Altmann, T., Stitt, M., Willmitzer, L., and Melchinger, A. E. (2012). Genomic and metabolic prediction of complex heterotic traits in hybrid maize. Nature genetics, 44(2), 217.

Rohart, F., San Cristobal, M., and Laurent, B. (2014). Selection of fixed effects in high dimensional linear mixed models using a multicycle ecm algorithm. Computational Statistics & Data Analysis, 80, 209–222.

Rubin, D. B. (1974). Estimating causal effects of treatments in randomized and nonrandomized studies. Journal of educational Psychology, 66(5), 688.

Schelldorfer, J., Bühlmann, P., and De Geer, S. V. (2011). Estimation for high-dimensional linear mixed-effects models using 1-penalization. Scandinavian Journal of Statistics, 38(2), 197–214.

Sohn, M. B., Li, H., et al. (2019). Compositional mediation analysis for microbiome studies. The Annals of Applied Statistics, 13(1), 661–681.

Sun, T. and Zhang, C.-H. (2010). Comments on: 1-penalization for mixture regression models. Test, 19(2), 270–275.

Sun, T. and Zhang, C.-H. (2012). Scaled sparse linear regression. Biometrika, 99(4), 879–898.

Tan, Z., Roche, K., Zhou, X., and Mukherjee, S. (2018). Scalable algorithms for learning high-dimensional linear mixed models. arXiv preprint arXiv:1803.04431.

Tian, J., Keller, M. P., Oler, A. T., Rabaglia, M. E., Schueler, K. L., Stapleton, D. S., Broman, A. T., Zhao, W., Kendziorski, C., Yandell, B. S., et al. (2015). Identification of the bile transporter slco1a6 as a candidate gene that broadly affects gene expression in mouse pancreatic islets. Genetics, pages genetics–115.

Tu, Z., Keller, M. P., Zhang, C., Rabaglia, M. E., Greenawalt, D. M., Yang, X., Wang, I.-M., Dai, H., Bruss, M. D., Lum, P. Y., et al. (2012). Integrative analysis of a cross-loci regulation network identifies app as a gene regulating insulin secretion from pancreatic islets. PLoS genetics, 8(12), e1003107.

VanderWeele, T. (2015). Explanation in causal inference: methods for mediation and interaction. Oxford University Press.

VanderWeele, T. and Vansteelandt, S. (2014). Mediation analysis with multiple mediators. Epidemiologic methods, 2(1), 95–115.

VanderWeele, T. J. (2011). Controlled direct and mediated effects: definition, identification and bounds. Scandinavian Journal of Statistics, 38(3), 551–563.

VanRaden, P. M. (2008). Efficient methods to compute genomic predictions. Journal of dairy science, 91(11), 4414–4423.

Wang, P., Dawson, J. A., Keller, M. P., Yandell, B. S., Thornberry, N. A., Zhang, B. B., Wang, I.-M., Schadt, E. E., Attie, A. D., and Kendziorski, C. (2011). A model selection approach for expression quantitative trait loci (eqtl) mapping. Genetics, 187(2), 611–621.

Xu, P., Wang, T., Zhu, H., and Zhu, L. (2015). Double penalized h-likelihood for selection of fixed and random effects in mixed effects models. Statistics in Biosciences, 7(1), 108–128.

Yuille, A. L. and Rangarajan, A. (2003). The concave-convex procedure. Neural computation, 15(4), 915–936.

Zhang, H., Zheng, Y., Zhang, Z., Gao, T., Joyce, B., Yoon, G., Zhang, W., Schwartz, J., Just, A., Colicino, E., et al. (2016). Estimating and testing high-dimensional mediation effects in epigenetic studies. Bioinformatics, 32(20), 3150–3154.

Zhao, Y., Lindquist, M. A., and Caffo, B. S. (2018). Sparse principal component based high-dimensional mediation analysis. arXiv preprint arXiv:1806.06118.

Zhu, Z., Zhang, F., Hu, H., Bakshi, A., Robinson, M. R., Powell, J. E., Montgomery, G. W., Goddard, M. E., Wray, N. R., Visscher, P. M., et al. (2016). Integration of summary data from gwas and eqtl studies predicts complex trait gene targets. Nature genetics, 48(5), 481.

Zou, H. (2006). The adaptive lasso and its oracle properties. Journal of the American statistical association, 101(476), 1418–1429.

## References

Jianan Tian, Mark P Keller, Angie T Oler, Mary E Rabaglia, Kathryn L Schueler, Donald S Stapleton, Aimee Teo Broman, Wen Zhao, Christina Kendziorski, Brian S Yandell, et al. Identification of the bile transporter slco1a6 as a candidate gene that broadly affects gene expression in mouse pancreatic islets. Genetics, pages genetics–115, 2015.

Zhidong Tu, Mark P Keller, Chunsheng Zhang, Mary E Rabaglia, Danielle M Greenawalt, Xia Yang, I-Ming Wang, Hongyue Dai, Matthew D Bruss, Pek Y Lum, et al. Integrative analysis of a cross-loci regulation network identifies app as a gene regulating insulin secretion from pancreatic islets. PLoS genetics, 8(12):e1003107, 2012.

Ping Wang, John A Dawson, Mark P Keller, Brian S Yandell, Nancy A Thornberry, Bei B Zhang, I-Ming Wang, Eric E Schadt, Alan D Attie, and Christina Kendziorski. A model selection approach for expression quantitative trait loci (eqtl) mapping. Genetics, 187(2):611–621, 2011.

